# High intensity muscle stimulation activates a systemic Nrf2-mediated redox stress response

**DOI:** 10.1101/2021.04.12.439469

**Authors:** Ethan L. Ostrom, Ana P. Valencia, David J. Marcinek, Tinna Traustadóttir

## Abstract

**Introduction:** High intensity exercise is an increasingly popular mode of exercise to elicit similar or greater adaptive responses compared to traditional moderate intensity continuous exercise. However, the molecular mechanisms underlying these adaptive responses are still unclear. The purpose of this pilot study was to compare high and low intensity contractile stimulus on the Nrf2-mediated redox stress response in mouse skeletal muscle.

**Methods:** An intra-animal design was used to control for variations in individual responses to muscle stimulation by using a stimulated limb (STIM) and comparing to the contralateral unstimulated control limb (CON). High Intensity (HI – 100Hz), Low Intensity (LI – 50Hz), and Naïve Control (NC – Mock stimulation vs CON) groups were used to compare these effects on Nrf2-ARE binding, Keap1 protein content, and downstream gene and protein expression of Nrf2 target genes.

**Results:** Muscle stimulation significantly increased Nrf2-ARE binding in LI-STIM compared to LI-CON (p = 0.0098), while Nrf2-ARE binding was elevated in both HI-CON and HI-STIM compared to NC (p = 0.0007). The Nrf2-ARE results were mirrored in the downregulation of Keap1, where Keap1 expression in HI-CON and HI-STIM were both significantly lower than NC (p = 0.008) and decreased in LI-STIM compared to LI-CON (p = 0.015). In addition, stimulation increased NQO1 protein compared to contralateral control regardless of stimulation intensity (p = 0.019).

**Conclusions:** Taken together, these data suggest a systemic redox signaling exerkine is activating Nrf2-ARE binding and is intensity gated, where Nrf2-ARE activation in contralateral control limbs were only seen in the HI group. Other research in exercise induced Nrf2 signaling support the general finding that Nrf2 is activated in peripheral tissues in response to exercise, however the specific exerkine responsible for the systemic signaling effects is not known. Future work should aim to delineate these redox sensitive systemic signaling mechanisms.

## INTRODUCTION

Exercise induces beneficial adaptations through redox signaling cascades that are mediated by the redox stress response transcription factor nuclear erythroid related factor 2 (Nrf2) [1-3]. The redox stress response system is a critical cytoprotective mechanism that protects both the cell from endogenous and environmental redox stressors and contributes to adaptive processes to exercise [4, 5]. The redox signal transduction cascade is a highly complex and coordinated system involving the generation of reactive oxygen species, the oxidation of redox-relay molecules or direct oxidation of sensor molecules with thiol switches like Kelch-like ECH-associated protein 1 (Keap1), and effector molecules that change activity, localization, protein-protein interactions, or protein turnover in response to the redox signaling cascade [6, 7]. The overall physiological adaptation resulting from redox signaling cascades depends on the rate of accumulation of these reactive species, and the steady-state levels of enzymatic and non-enzymatic antioxidants [4], as well as the basal oxidation state of proteins with thiol-based redox switches [7-9].

Reactive oxygen species (ROS) generation is required for appropriate skeletal muscle adaptations to exercise such as mitochondrial biogenesis [10]. Inhibiting exercise-induced ROS signaling with antioxidants impairs markers of mitochondrial biogenesis including AMPK and PGC1α activation [11-13], Nrf2 activation [2], and downstream gene expression [14]. Furthermore, there is a linear relationship between exercise-induced ROS accumulation and Nrf2 activation [15], which is dependent on the duration of the exercise bout [15, 16]. Nrf2 can also be activated in human skeletal muscle in response to supramaximal exercise of shorter duration [17], suggesting that longer durations are not required to induce Nrf2 activity if the intensity of the bout is high enough. High intensity interval training is a popular training method to obtain improvements in cardiorespiratory fitness quickly and efficiently, partly through increases in mitochondrial biogenesis – processes that are redox signaling dependent [10, 18-20].

The molecular pattern of redox signaling responses can differ depending on the intensity of the exercise which leads to different adaptations and resulting phenotypes [21-23]. However only two studies to date have directly compared moderate intensity and high intensity exercise and Nrf2 activation, one in human peripheral blood mononuclear cells (PBMCs), and one in human skeletal muscle [24, 25]. There were no differences in Nrf2 signaling between exercise intensities in either human PBMCs or human muscle [24, 25]. However other studies comparing different exercise intensities have shown divergent redox signaling effects in other redox proteins, highlighting some interesting caveats in the field [26]. Therefore, more detailed and well controlled studies comparing differing intensity stimuli on Nrf2 redox stress response signaling are needed.

One issue regarding the redox stress response to exercise, is the considerable variability in the adaptive signaling responses across individuals [27]. This makes it difficult to predict the hormetic response, and thus the overall adaptation to regular exercise, and impacts the ability to effectively prescribe exercise to clinical and non-clinical populations. Therefore, the purpose of this pilot study was to test high and low muscle stimulation intensities on redox stress responses in mouse skeletal muscle using a within-animal study design to control for any intra-animal variation in responses to the contractile stimulus, as well as a between-animal control using a mock stimulation condition. We hypothesized that high intensity stimulation would lead to greater adaptations and activation of Nrf2 signaling compared to low intensity exercise. Here we show a novel systemic Nrf2-ARE activation mechanism gated by high intensity stimulation but not low intensity stimulation that increases Nrf2-ARE binding by acting through inhibition of the negative regulator Keap1.

## METHODS

### Animals

Young (6mo) C57BL/6 male mice were received from the Jackson Labs. All mice were maintained at 21°C on a 14/10 light/dark cycle and given standard mouse chow and water *ad libitum*. The study was approved by the University of Washington Institutional Animal Care and Use Committee (IACUC) and the tissue transfer was approved by the Northern Arizona University IACUC.

### Study Design

There were three experimental groups for this study (Figure 1): High intensity stimulus (HI, n=5), low intensity stimulus (LI, n=5), and naïve unstimulated control (NC, n=5). An intra-animal study design was used for HI and LI groups to control for within animal variation, where the stimulated (STIM) limb was used as the exercise condition and contralateral unstimulated (CON) limb was used as the non-exercise control. The Naïve Controls (NC) were used as the inter-animal controls and were anesthetized for the same amount of time as the other two groups before both hind limbs were harvested, but no muscle stimulation occurred in either limb. Naïve controls were used to control for systemic effects that could affect both limbs in the HI and LI groups. Nrf2-mediated redox stress response signaling was compared between each group (HI, LI, & NC) as well as paired comparisons within animals unstimulated control and stimulated (exercised) limbs (Figure 1).

**Figure 1.**
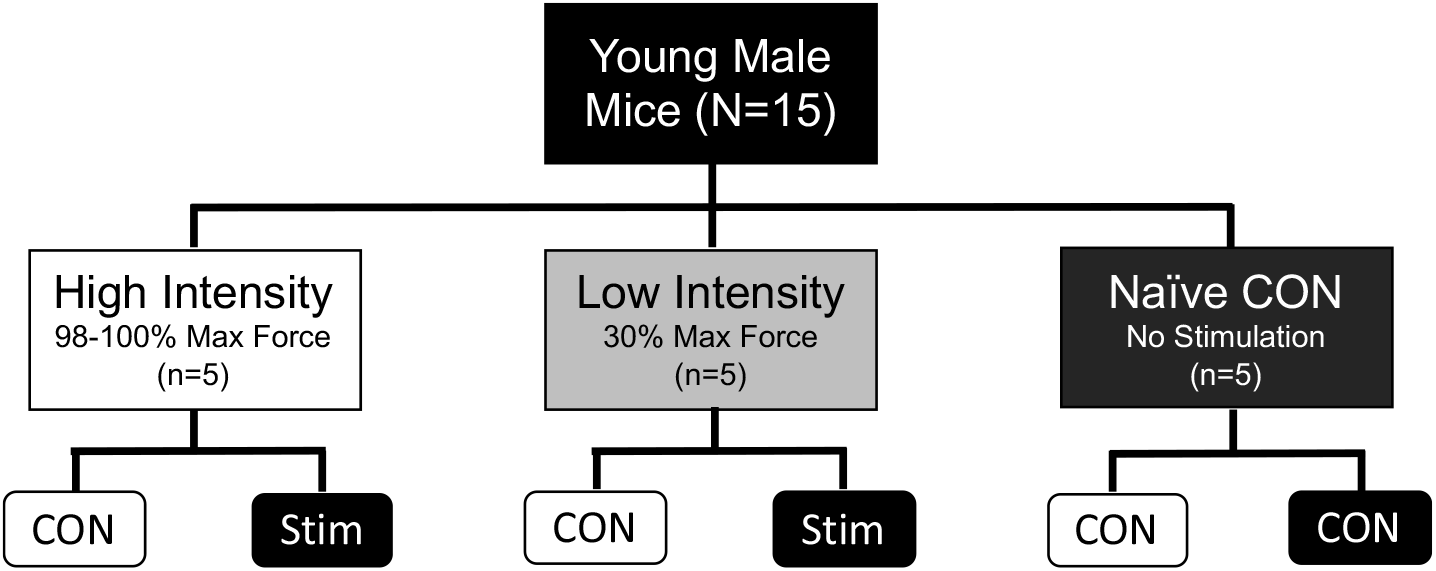
Study Design

### In Vivo Muscle Stimulation

Muscle stimulation was performed on the right leg of anesthetized mice (1-2% isofluorane) resting on a heated plate at 37°C as previously described [28]. The knee was secured, and foot taped to a footplate perpendicular to the tibia. The footplate was connected to a force transducer (Aurora Scientific, ON, Canada). The tibial nerve was stimulated using Grass Stimulator (S88X, Astro Med, Inc.) at optimal voltage (1-4V) that was selected by measuring maximal isometric torque of plantarflexion during isometric contractions (200ms train, 0.1ms pulse, 100Hz). Following optimization, the plantarflexors underwent a fatigue protocol with isometric contractions (200ms train, 0.1ms pulse) induced by a high (100Hz) or low (50Hz) stimulation frequency every fourth second for 30 minutes. Following the fatigue procedure, the electrodes were removed and the was leg released from the instrument. While on anesthesia, mouse remained on the heated plate until muscle harvest. The gastrocnemius and soleus muscle from the stimulated and unstimulated legs (contralateral control) were removed 30 minutes after the end of muscle stimulation and flash frozen in liquid nitrogen and stored at -80°C. Animals were euthanized through cervical dislocation.

### Gene Expression

Soleus muscles were homogenized in RLT buffer (+ 1% BME). RNA extraction was done using RNeasy Plus Mini Kit following Proteinase K digestion and elimination of DNases using RNase-free DNase kit (all reagents from Qiagen). Isolated RNA was then converted to cDNA using Bio-Rad iScript kit and RT-qPCR using Bio-Rad SYBR Sso Advanced. Samples were analyzed using the ΔΔCt method. A panel of six housekeeping genes were used to determine which were the best three genes to use for internal controls (Table 1). The geometric mean for the three most stable housekeeping genes were used to quantify changes in target gene mRNA expression in response to stimulation [29]. All primer sequences are listed in Table 1.

**Table 1.**
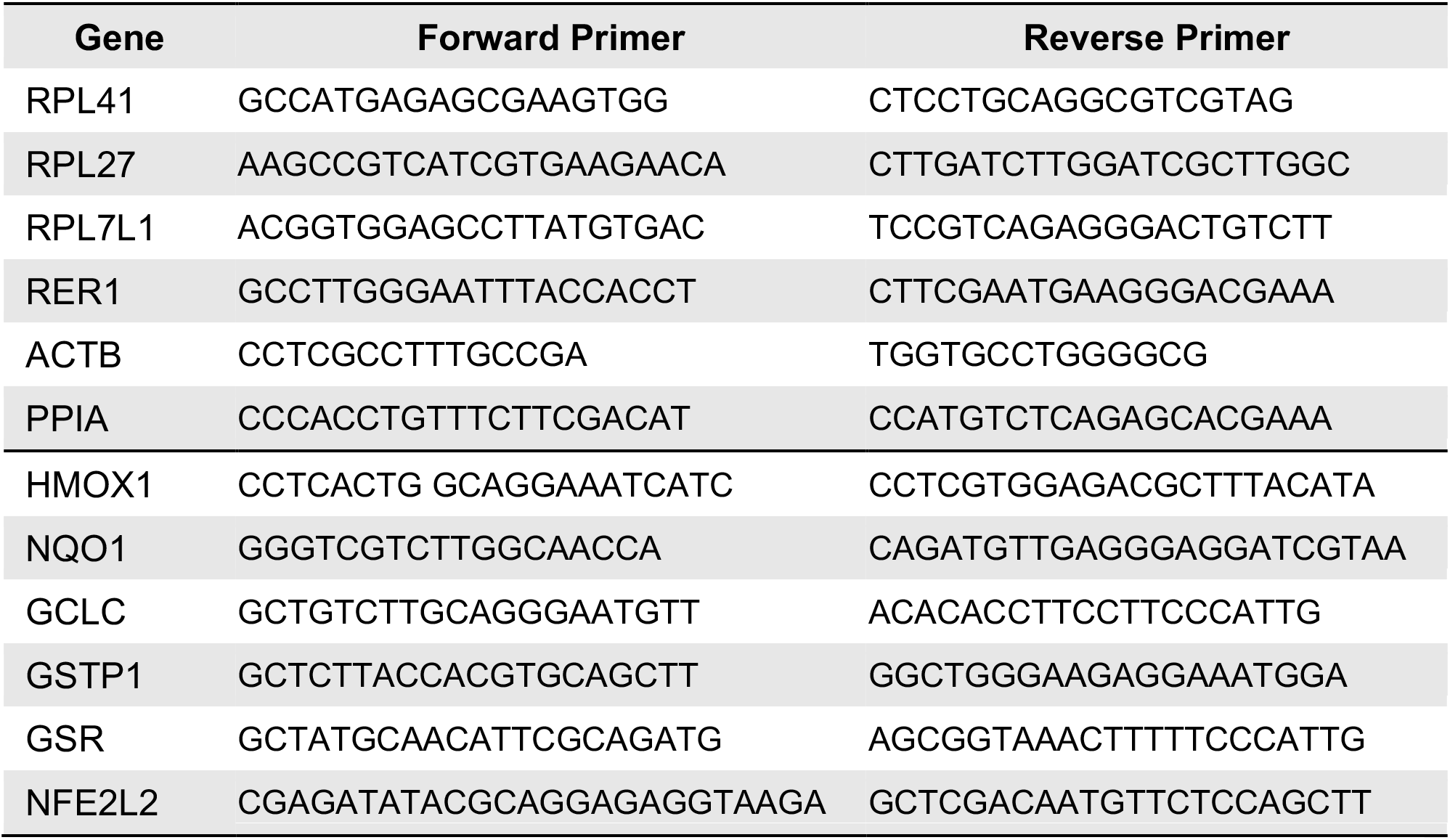
Primer Sequences for RT -qPCR.

### Glutathione Content

Intracellular glutathione levels were measured as previously described [30] using a spectrophotometric assay based on the affinity of 2,3-naphthalenedicarboxaldehyde (NDA) for γ-glutamylcysteinylglycine (GSH). Skeletal muscle (10-20mg) was homogenized in ice-cold Locke’s buffer (10 mM HEPES, 5.5 mM KCl, 10 mM glucose, 5 mM NaHCO3, and 130 mM NaCl). A portion of the homogenate was set aside for protein quantification using a Bradford assay. An equal volume of 200mM 5-sulfosalicylic acid dehydrate (SSA) was mixed in. After resting on ice for 15 minutes the samples were centrifuged at 12,000 x g for 3 minutes at 4 °C. The supernatant was plated into a 96 well plate in duplicate. Standards with known GSH concentrations (0-50µM) were also plated in duplicate. 0.2 M NEM/0.02 M KOH were added to each well followed by 10 mM TCEP. After a 20-minute incubation at room temperature, 0.5 N NaOH was added followed by 10mM NDA. After a 30-minute incubation the plate was read at fluorescence intensity using 472 (excitation) and 528 (emission). GSH levels were assessed using the standard curve and normalized to protein concentration.

### Western Blotting

Gastrocnemius muscles were homogenized on ice for 5 minutes in the presence of 25µl of Cell Lytic MT cell lysis buffer (Sigma-Aldrich) per mg of tissue. Lysis buffer was supplemented with 0.1% protease inhibitors (Sigma-Aldrich) and 0.2% phosphatase inhibitor (ThermoFisher). Following homogenization, aliquots were taken from original samples for Bradford assays to determine protein concentrations. Next, sample buffer was added to the original sample then boiled and stored at -80°C. Thirty micrograms of protein were separated by gel electrophoresis in each well followed by wet transfer on to nitrocellulose membranes and one hour blocking in TBS + 5% non-fat dry milk. Blots were incubated with Keap1 (EPR22664-26, Abcam, Cambridge MA) GCLC (EP12345, Abcam), GSR (Santa Cruz Biotech, Dallas TX), HO1 (Cell Signaling Technology, Danvers MA), or NQO1 (Cell Signaling Technology, Danvers MA) monoclonal antibodies to detect proteins with Ponceau S, used for loading controls (Cell Signaling Technology, Danvers MA).

### Nrf2-ARE Binding Assay

Gastrocnemius muscles were homogenized, lysed, and nuclear fractions extracted per manufacturer’s instructions (Nrf2-ARE binding kit, TransAM, Carlsbad CA). Briefly, samples were homogenized in a 1:30 ratio (mg tissue:µl buffer) followed by centrifugation, isolation, and lysis of the nuclear pellet. After Bradford assays were run to determine protein content in nuclear fractions, nuclear lysates were incubated in a 96 well plate coated with ARE consensus oligonucleotides for Nrf2 Binding. Following three washes, primary Anti-Nrf2 antibodies were incubated (1:1000) in each well used to detect Nrf2 followed by secondary antibody incubation and colorimetric development. Absorbance was read at 450nm on a Synergy HT plate reader (Bio-Tek, Winooski, VT).

### Statistical Analysis

For all measures, a three by two repeated measures ANOVA (Intensity group by Stimulation condition) was run for analysis of intensity group differences, stimulation differences and their interaction. Tukey’s post hoc analysis was used for CON vs STIM pairwise comparisons within group. All relevant p values reported are adjusted p values using Tukey’s multiple comparisons post hoc analysis. Statistical analysis was performed using SPSS (IBM, version 27) and GraphPad Prism software (San Diego, CA).

## RESULTS

### Muscle stimulation

The results from the muscle stimulation protocol in HI and LI are shown in figure 2. The force output of a tetanic contraction at 100Hz was not different between groups (12.1 mN-m ± 0.9 and 11.9 mN-m ± 1.7 for LI-STIM and HI-STIM respectively). The fatigue protocol at HI-STIM led to a final force output that was 27.7% ± 5.6 of initial 100Hz tetanus, and LI-STIM led to a final force output that was 12.8% ± 5.6 of a 100Hz tetanus and 37.3% ± 2.8 of the initial 50Hz contraction.

**Figure 2.**
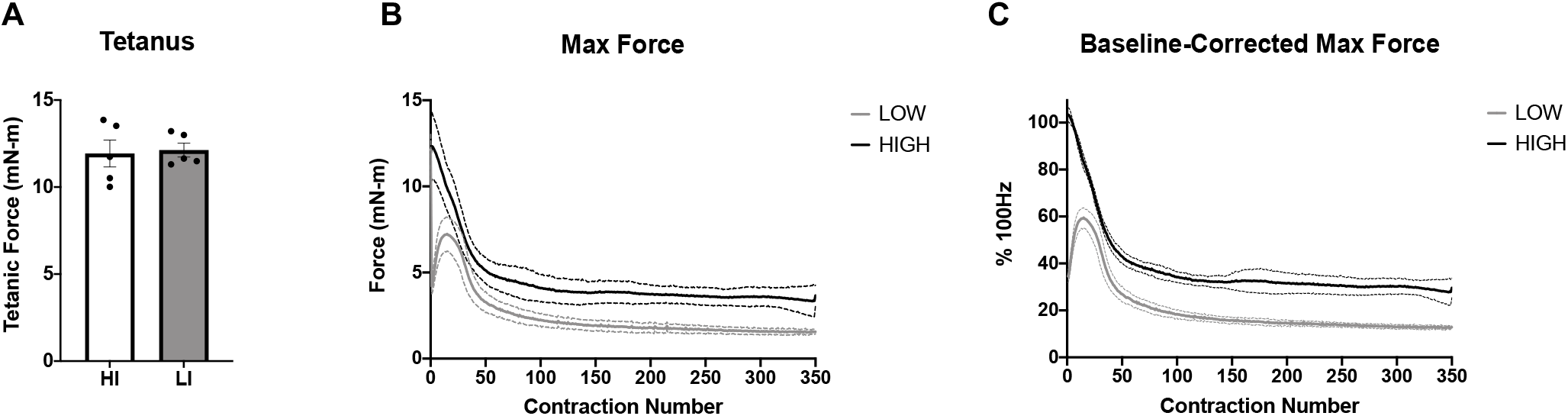
Muscle force output and fatiguing stimulation. A) Both low and high intensity stimulation groups had similar maximal tetanic force at 100Hz. B) The absolute force (mN-m) over time in response to either High (100Hz) or Low (50Hz) intensity stimulation frequency. C) The normalized force to maximal tetanus. Data are Mean ± SD.

### Stimulation Intensity Reveals Divergent Effects on Nrf2-ARE Binding and Keap1 Protein Dynamics in Skeletal Muscle

Nrf2-ARE binding was low under basal unstimulated conditions (Figure 3A, Naïve Control group and LI-CON condition). There was a significant interaction between stimulation condition by intensity (RM ANOVA Group x Stimulation condition, p = 0.005), where Nrf2 ARE binding significantly increased in LI-STIM condition compared to LI-CON (Figure 3A, ** p = 0.047), while NC and HI were not significantly different between conditions (Figure 3A, HI-CON vs HI-STIM, or both NC-CON conditions). Nrf2-ARE binding was significantly higher in the HI group under both conditions compared to NC group (Figure 3A, ^##^ p = 0.007). Keap1 showed a significant main effect of intensity indicated by the difference between high intensity and naïve controls (Figure 3B, ^##^p = 0.008). There was also an interaction of stimulation by intensity (p = 0.035) and a main effect of stimulation (p = 0.016). The interaction of stimulation by intensity was driven by a significant decrease in Keap1 protein in LI-STIM condition compared to LI-CON (Figure 3B, *p = 0.015) with no changes between CON vs STIM conditions in HI or differences between the two NC-CON. There were no differences in total glutathione content across groups, or in response to stimulation within groups (Figure 3D).

**Figure 3.**
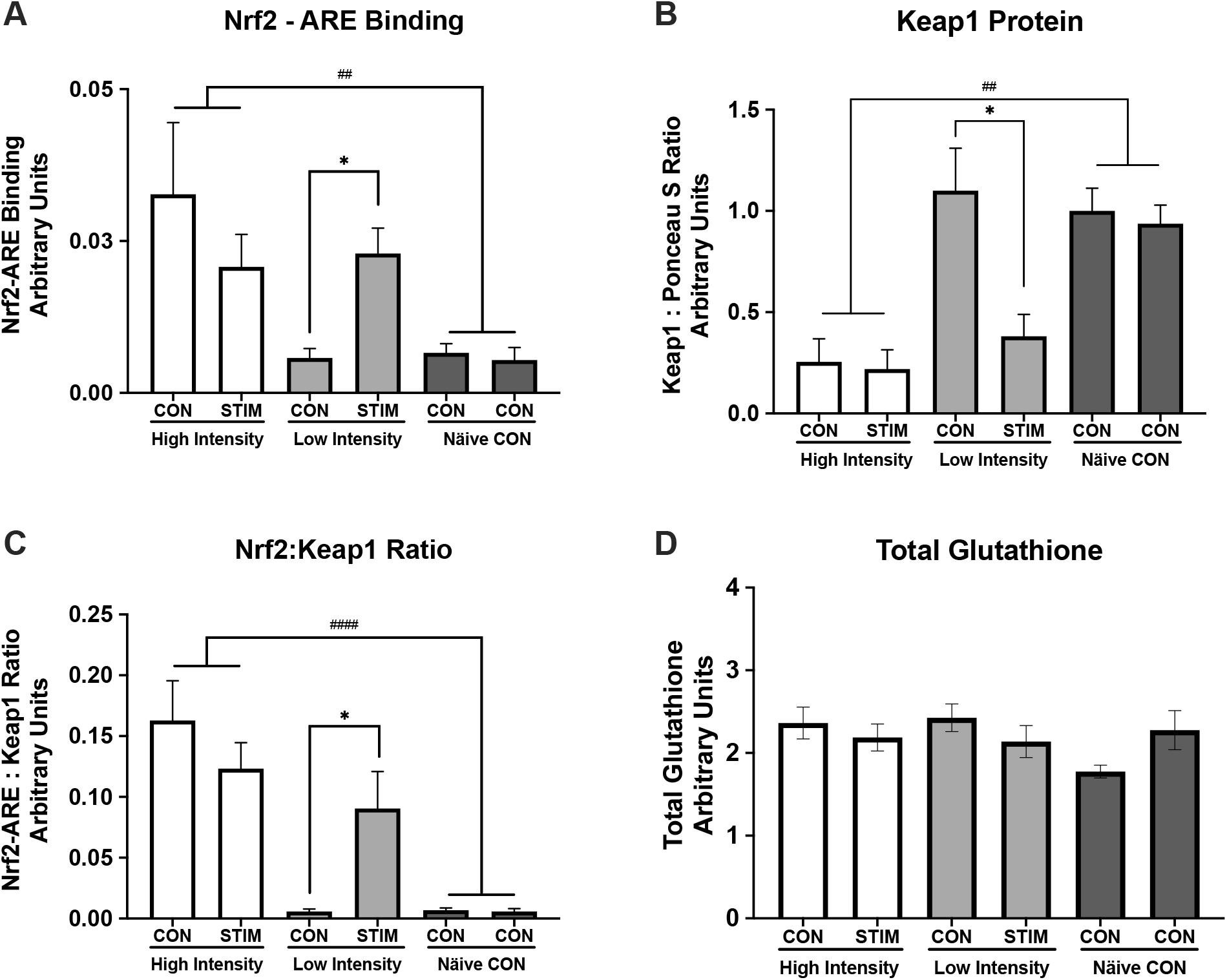
Nrf2-ARE Binding and Keap1 Protein Expression in Response to Skeletal Muscle Stimulation. Keap1-Nrf2-ARE signaling response to high and low intensity stimulation and total glutathione content. A) Nrf2-ARE binding is low under basal unstimulated conditions (Naïve Control group and LI CON condition) but increases in response to low intensity stimulation (LI-CON vs LI-STIM, * p = 0.047). Nrf2-ARE binding is significantly higher in the HI group compared to NC group (^##^ p = 0.007) . B) Keap1 content is unchanged between limbs in the Naïve Control group or the LI-CON, however there is a significant decrease in Keap1 content in the LI-STIM limb (*p = 0.015). Keap1 content is significantly lower in the HI group under both conditions compared to NC (^##^p = 0.008). C) Nrf2:Keap1 ratio illustrates the same pattern as either Nrf2-ARE binding or Keap1 protein content alone (HI vs NC, ^####^ p < 0.0001). D) Total glutathione content was not different across groups and did not change significantly in response to stimulation. Representative western blot image of Keap1 protein is shown in Figure 5B. Keap1 was normalized to Left hind limb of the NC group. Data are presented as mean ± SEM.

### Gene expression responses to skeletal muscle stimulation

A panel of six housekeeping genes were used to assess their stability in response to skeletal muscle stimulation (Figure 4A and 4B). Variability from CON to STIM conditions was assessed for each housekeeping gene (Figure 4A) as well as variation by intensity group (Figure 4B). RPL41, RPL27, and RPL7l1 were selected as the best genes to use for analysis because their within animal variation was lowest across groups (ΔCq in response to stimulation was minimized). Target gene fold change was calculated using the geometric mean of the 3 housekeeping genes with the ΔΔCq method. Target gene responses to muscle stimulation were not significantly different from CON or across intensities, although there was a trend for an increase in GCLC mRNA expression in response to stimulation (Figure 4C, p = 0.06).

**Figure 4.**
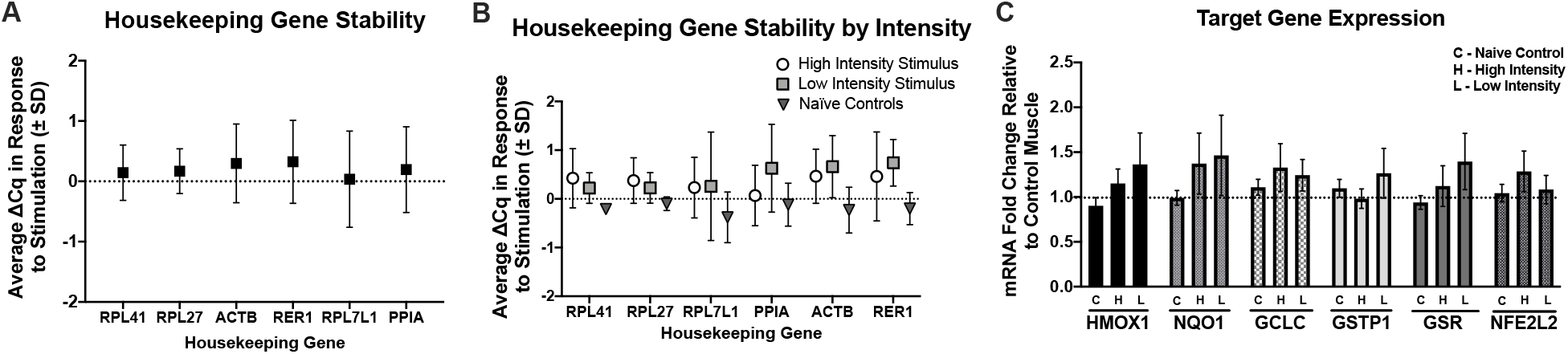
Gene Expression of Redox Stress Response Genes to Skeletal Muscle Stimulation. A) Housekeeping gene average Cq change (Δ Cq) in response to skeletal muscle stimulation. B) Housekeeping gene average Cq change (Δ Cq) by group. Based on these results RPL41, RPL27, and RPL7l1 were selected for housekeeping genes. C) Effects of high intensity (H), low intensity (L) muscle stimulations, and control (C) for target genes HMOX1, NQO1, GCLC, GSR, and NFE2L2 fold change from unstimulated muscle (dotted line). In Naïve Controls the mock stimulated muscle (right limb) was compared to the unstimulated control (left limb in all cases). There were no statistically significant increases in gene expression in response to stimulation in either group, although GCLC showed trends for increases regardless of intensity groups. RPL41, RPL27, and RPL7l1 were used for analysis of target gene fold change. H = High intensity stimulation group, L = Low intensity stimulation group, C = Naïve control group.

### Changes in redox stress response proteins in response to skeletal muscle stimulation

There was a main effect of intensity for HO1 Protein expression (p = 0.01) with HO1 significantly elevated in the high intensity group compared to the Naïve controls (Figure 5C, p = 0.003). There was a significant effect of stimulation condition on NQO1 protein (Figure 5D, p = 0.019) with no differences between high and low intensity stimulus and no differences between stimulation groups. There was a significant main effect of stimulation on GCLC protein content where the stimulation condition decreased GCLC protein slightly (Figure 5E, p = 0.03) with no differences between groups. There were no significant differences between groups or in response to muscle stimulation for GR protein (Figure 5F).

**Figure 5.**
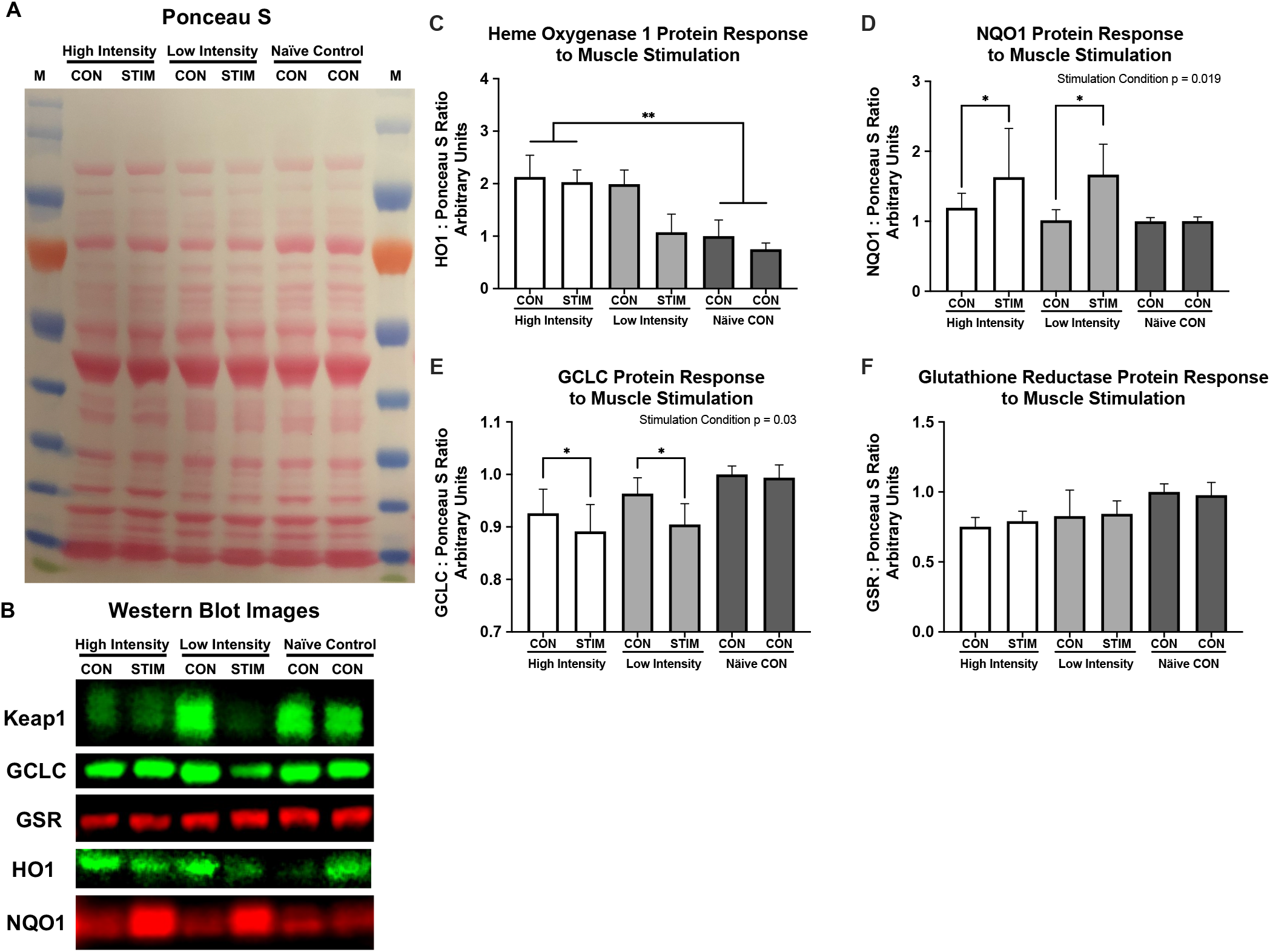
Redox stress protein response to muscle stimulation. Redox stress protein response to muscle stimulation. A) Representative Ponceau S stain, B) Representative western blot images. C) Heme Oxygenase 1 protein is significantly elevated in both conditions of the high intensity group compared to the Naïve control group (p = 0.002). D) NQO1 protein increased significantly in response to stimulation (Stimulation condition p = 0.019), with no differences between intensities, or across groups. E) GCLC decreases slightly in stimulation conditions regardless of intensity (Stimulation condition p = 0.03) with no differences between groups. Glutathione reductase is unchanged in response to muscle stimulation or across intensity groups. All values are normalized to left limb of the NC group and set equal to 1 for western blot graphs.

## DISCUSSION

Exercise is one of the most powerful pleiotropic interventions to improve health and fitness. These health benefits are mediated in part by redox signaling responses to an exercise bout, including activation of the inducible redox stress response transcription factor Nrf2. Nrf2 mediates hundreds of different cytoprotective and metabolic genes increasing metabolic and redox capacity with repeated transient stressors like exercise training. Recently there has been an increased interest in using high intensity interval exercise as a more time efficient way to elicit beneficial adaptations [18]. How the molecular signature from high intensity exercise differs from moderate intensity exercise is still unclear. The aim of this pilot study was to investigate the effects of high intensity and low intensity muscle stimulation on redox stress response markers in mouse skeletal muscle.

Previous studies on Nrf2 signaling responses to exercise have used whole animal treadmill exercise, measuring Nrf2 in several different tissues [2, 15, 16, 31-37], or whole-body exercise in humans [1, 17, 24, 25, 38-40]. We have recently shown that basal levels of Nrf2 affect its inducibility to an acute exercise bout in humans [39]. Therefore, in order to control for intra-animal variations in basal redox homeostasis and basal Nrf2 activation, we elected to use an intra-animal design, where the right limb of the animals were stimulated with either high intensity or low intensity muscle stimulation, while the contralateral unstimulated limb served as the internal control. Unexpectedly, we found that both stimulated and contralateral unstimulated limb of the high intensity group showed increased Nrf2-ARE binding activity. This is in contrast to the low intensity group where Nrf2 was low in the unstimulated contralateral control limb, but significantly increased in the stimulated limb. The low levels of Nrf2-ARE binding in both Naïve control limbs confirm that Nrf2 activity is low in resting healthy tissue.

Western blots of Keap1, the negative regulator of Nrf2, show opposite effects, where Keap1 protein is highly expressed in both naïve control limbs, and the low intensity control limb, as predicted with the low levels of Nrf2-ARE binding. Keap1 decreases in response to low intensity stimulation, releasing the inhibition on Nrf2 and resulting in increase in Nrf2 activation and ARE binding in low intensity stimulation. Keap1 is lowly expressed in both control and stimulated limbs of the high intensity group, in line with the increases in Nrf2 activation in both stimulated and contralateral control limbs. The ratio of Nrf2-ARE binding to Keap1 content also illustrates the relationship between Keap1 protein and Nrf2-ARE binding. A recently published paper reported significant decreases in Keap1 content after acute exhaustive exercise in human skeletal muscle and concomitant increases in Nrf2 [17], our data here are in agreement with these results. Together these data demonstrate i) Basal Nrf2 activation is low under resting or unstressed conditions in young adult skeletal muscle; ii) Nrf2 activity is inducible in response to muscle stimulation; iii) high intensity, but not low intensity muscle contraction, activates Nrf2 even in unstimulated muscles; and iv) Keap1 protein content mirrors these findings, indicating these inducible Nrf2 responses are acting through canonical Keap1 inhibition in response to skeletal muscle stimulation.

We measured gene expression 30 minutes after the muscle stimulation was finished. The timing may have been a limitation in detecting peak changes in gene expression of these redox stress response genes because there were no statistically significant increases. We and others have shown Nrf2 regulated genes to be induced between 1-4 hours after completion of an exercise bout [38, 39]. However, these results were in human PBMCs, and given that NQO1 protein content increased in response to stimulation in the current investigation, the gene expression may occur earlier in skeletal muscle than in PBMCs. Therefore, future research will utilize different timepoints to assess gene expression changes.

Our results show minimal changes to GSR, and slight but significant decreases in GCLC protein in response to muscle stimulation. The decrease in GCLC here is likely a temporal effect, where the acute stimulation increases proteasomal activity causing decreases in constitutively expressed proteins like GCLC, and the increase in protein likely occurs after the increase in mRNA expression. We have shown that GCLC protein increases significantly eight hours after a single bout of exercise, which is dependent on the increases in mRNA at 1 and 4 hours after completion of the exercise bout [39]. Therefore, increasing the time of skeletal muscle harvest would likely capture greater GCLC protein accretion in response to the exercise bout.

In contrast to GCLC protein expression, HO1 and NQO1 protein increased, albeit in different patterns, in response to muscle contraction. The elevated HO1 expression in both limbs of the high intensity group mirrors the activation of Nrf2 and suggests that HO1 protein expression is rapidly increased and intensity dependent. HO1 mRNA induction has also been shown to be intensity or dose dependent in human skeletal muscle, supporting these findings [41]. NQO1 protein was inducible in response to muscle stimulation in both high and low intensity muscle stimulation, with no significant differences between groups. NQO1 induction in response to exercise has been demonstrated in other reports [42] and is highly dependent on Nrf2 induction [43]. The fact that NQO1 mRNA was not statistically significant but the protein was, suggests that the time course for NQO1 mRNA accretion and protein is shifted closer to the exercise bout than GCLC mRNA. This poses an interesting paradox where some redox stress response genes are early response genes, while others may be late response genes. This may be due to other inducible transcription factors and inhibitors differentially regulating phase II antioxidant genes. For example, NF-kB is known to also regulate GCLC/M gene transcripts [44] which could be responsible for the discrepancies between gene transcripts and protein accumulation in our results.

Taken together, these data suggest that there is an intensity gated redox sensitive exerkine released from high intensity contraction, but not low intensity contraction, that is driving these divergent Nrf2 signaling events in response to high and low intensity exercise, respectively. Recent work has shown that pH gated release of succinate is a myokine involved in adaptive responses to exercise [45]. Others have shown that peroxiredoxins and thioredoxins are released into the blood plasma compartment in an intensity and time dependent manner in response to high intensity, but not moderate intensity exercise in humans [26]. While we cannot ascertain what exerkine is involved with our current results, the literature suggest that intensity gating mechanisms may explain the divergent effects seen in molecular responses to exercise, and that these mechanisms may be redox dependent [24, 26, 46]. This intensity gated systemic redox signaling model is illustrated in Figure 6, where the signal is released into the blood stream in response to the high intensity exercise and activates Nrf2 signaling in the contralateral unstimulated muscle, while low intensity muscle stimulation does not meet the threshold to propagate a systemic redox signaling response.

**Figure 6.**
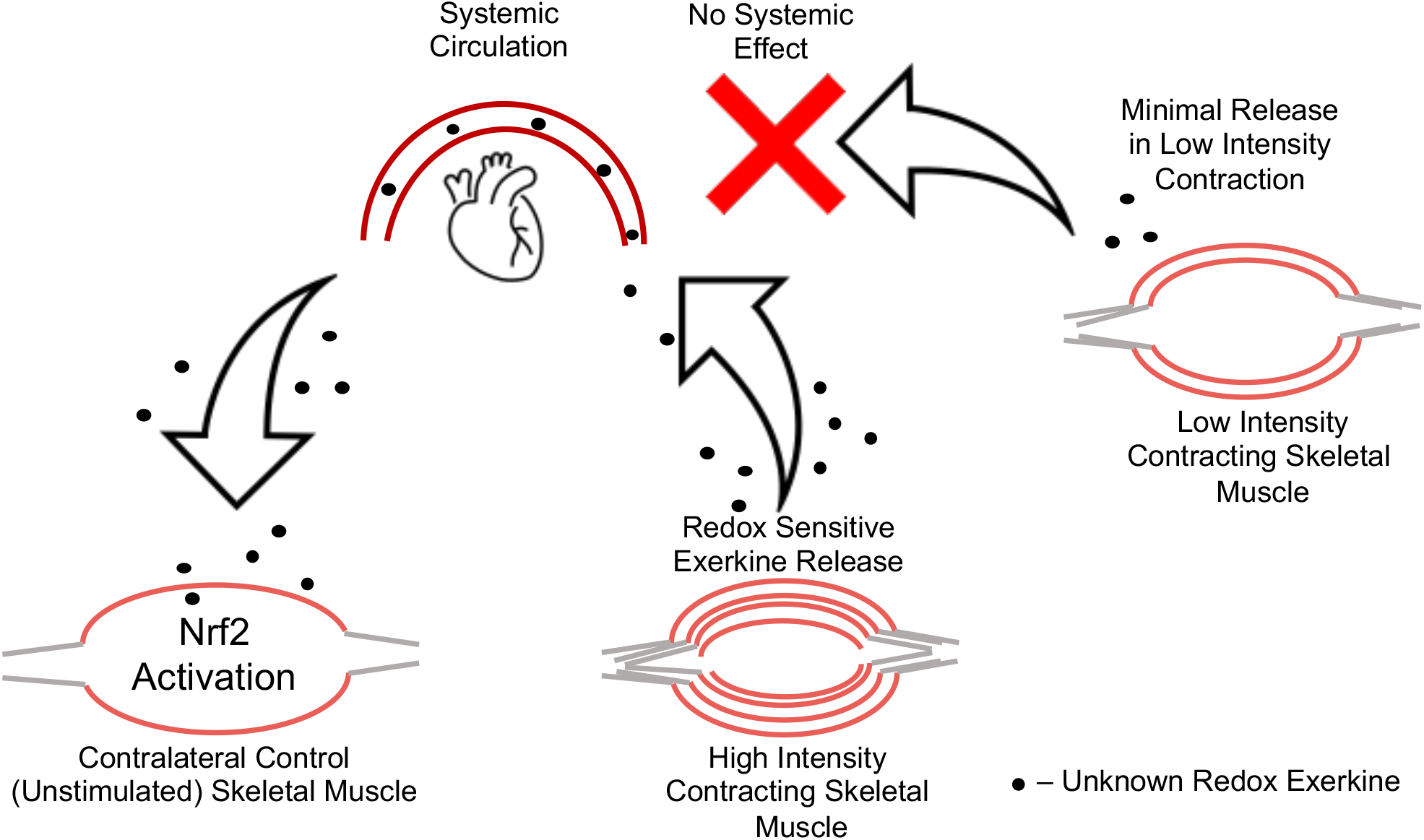
Model of systemic redox exerkine signaling is gated by muscle stimulation intensity. High intensity muscle contraction induces release of a redox active exerkine that travels through the blood stream and acts on unstimulated skeletal muscle. This effect is not seen in low intensity contralateral control muscle, suggesting the redox exerkine release from skeletal muscle is intensity gated. In other words, this redox exerkine is only released upon stimuli above a certain stress threshold or is released at lower intensities but not at sufficient concentrations to cause signaling effects in other tissues.

### Strengths, Limitations, and Future Directions

The strength of the current study is the utilization of multiple controls to assess skeletal muscle redox signaling: Internal contralateral unstimulated control limbs compared to stimulated limbs, as well as Naïve Control animals with control limbs and “mock” stimulated control limbs. However, one limitation is the redox balance measures in this cohort. Total glutathione was measured, but after partitioning the sample for each assay there was not enough left to perform any additional measures of NAD^+^/NADH ratio, ROS production assays, or other redox balance measures. While additional redox balance markers would provide a more complete picture, perhaps more relevant to the current study is thiol redox status in target proteins. Future work should investigate protein thiol redox state in key signaling proteins like Keap1 in response to exercise. It is well known that Keap1 thiols are oxidized in response to oxidative stress, and that these modifications activate Nrf2. However, we are unaware of any study that has directly investigated the redox status of Keap1 protein thiols in response to exercise specifically. Furthermore, in studies measuring Nrf2 in tissues other than muscle tissue, it is still unclear how redox signaling mechanisms are being propagated.

Future investigations should aim to identify the redox exerkine responsible for systemic signaling effects in the inducible redox stress response signaling system. In these studies that demonstrate exercise induced Nrf2 activation in peripheral tissues like nervous tissue, PBMCs, and lung tissue, and the current investigation, it is unlikely that superoxide and hydrogen peroxide are viable secondary messenger signaling candidates given their half-life and concentration of extracellular enzymes capable of quenching these signals. The prerequisite for this specific redox myokine is that it is stable enough to travel through the bloodstream and affect other peripheral tissues, including skeletal muscle. It is also possible that there is a relay of some kind, where the exerkine can bind to a surface receptor on a neighboring cell that initiates an intracellular signaling cascade which elicits ROS production. The sheer number of possible candidate-myokines including proteins, lipids, RNA, exosomes, and microRNAs that could be responsible for the peripheral redox signaling makes this a difficult task. These findings are consistent with previous literature on high intensity exercise eliciting different molecular signaling cascades than low intensity exercise and illustrate an interesting dichotomy between low intensity and high intensity exercise. However, this is the first report that we are aware of demonstrating Nrf2 activation in response to high intensity but not low intensity exercise in a contralateral non-exercised muscle. These findings provide exciting future directions to unveil the novel signaling mechanisms highlighted here.

## Conclusions

This pilot study set out to test the effects of muscle contraction intensity on Nrf2-mediated redox signaling. The current study design demonstrates a muscle contraction induced systemic redox signaling that appears to be intensity gated; where contralateral unstimulated muscles are responding to high intensity, but not low intensity stimulation. These effects are exerted in part on the Keap1-Nrf2-ARE signaling axis, as well as downstream gene and protein expression for the major Nrf2 targets: NQO1 and HO1 protein. While these data are still preliminary, follow-up experiments are being designed to confirm these findings, including discovering the type of molecule or set of molecules responsible for these systemic redox signaling effects.

